# Threat imminence dictates the role of the bed nucleus of the stria terminalis in contextual fear

**DOI:** 10.1101/696112

**Authors:** Travis D. Goode, Gillian M. Acca, Stephen Maren

**Author notes:** Corresponding author, Address: Stephen Maren, Department of Psychological and Brain Sciences, Texas A&M University, 301 Old Main Dr., College Station, TX 77843-3474.

## Abstract

Previous work indicates that the bed nucleus of the stria terminalis (BNST) is involved in defensive freezing to unpredictable Pavlovian conditioned stimuli (Goode et al., 2019). Here we show that the BNST mediates freezing to contexts paired with remote (unpredictable), but not imminent (predictable), footshock. Rats underwent a fear conditioning procedure in which a single footshock unconditioned stimulus (US) was delivered either 1 (imminent) or 9 minutes (remote) after placement in the context; each rat received a total of four conditioning trials over two days. Contexts associated with either imminent or remote USs produced distinct patterns of freezing and shock-induced activity but freezing in each case was context-dependent. Reversible inactivation of the BNST reduced the expression of contextual freezing in the context paired with remote, but not imminent, footshock. Implications of these data are discussed in light of recent conceptualizations of BNST function, as well as for anxiety behaviors.

## INTRODUCTION

Anxiety disorders, such as generalized anxiety disorder (GAD), social anxiety disorder, and panic disorder, as well as stress and trauma disorders, such as posttraumatic stress disorder (PTSD), are among the most common and debilitating of mental illnesses (Craske et al., 2017; Essau et al., 2018; Kilpatrick et al., 2013; McMillan et al., 2014; Ravindran and Stein, 2010; Stein et al., 2017). Despite their prevalence and severity, a complete understanding of the brain mechanisms of anxiety-related behaviors has been elusive. Current models indicate that anxiety and trauma disorders involve a complex network of highly interconnected brain regions (Adhikari, 2014; Avery et al., 2016; Brooks and Stein, 2015; Calhoon and Tye, 2015; Ch’ng et al., 2018; Dunsmoor and Paz, 2015; Fenster et al., 2018; Fox and Shackman, 2019; Janak and Tye, 2015; Lebow and Chen, 2016; Maren et al., 2013; Miles and Maren, 2019; Robinson et al., 2019; Shackman and Fox, 2016); these include (but are not limited to) the medial prefrontal cortex, amygdala, hippocampus, and bed nucleus of the stria terminalis (BNST). In recent years, growing interest has centered on the BNST as a potential target of therapeutic interventions. However, the precise circumstances that engage the BNST in the learning and memory processes involved in anxiety are poorly understood.

To address these lingering questions, we have used Pavlovian fear conditioning procedures to probe the contributions of the BNST to aversive learning and memory. In this form of learning, a neutral conditioned stimulus (CS), such as an auditory tone, is paired with a salient and aversive stimulus, such as footshock [unconditioned stimulus (US)] (Maren, 2001a; Pavlov, 1927; Rescorla, 1968, 1988). The US itself elicits a number of unconditioned behaviors (URs), including bursts in activity and ultrasonic vocalizations (Fanselow, 1994). With as little as a single pairing with the US, the CS alone will elicit conditioned defensive responses (CRs; including defensive immobility or “freezing”, which often serves as the index of conditioning). During fear conditioning, animals not only learn that the CS predicts the US, but also learn to fear the context in which conditioning occurs. Interestingly, numerous studies implicate the BNST in the acquisition and expression of conditioned fear to contexts, but not discrete CSs (Goode et al., 2015; LeDoux et al., 1988; Luyten et al., 2011; Poulos et al., 2010; Resstel et al., 2008; Sullivan et al., 2004; Waddell et al., 2006; Walker and Davis, 1997; Zimmerman and Maren, 2011).

A number of interpretations have been developed to explain the selective role for the BNST in contextual fear (Davis et al., 2010, 1997b, 1997a; Fox et al., 2015; Fox and Shackman, 2019; Gafford and Ressler, 2015; Goode and Maren, 2017; Gungor and Paré, 2016; Klumpers and Kroes, 2019; Luyck et al., 2019; Miles and Maren, 2019; Robinson et al., 2019; Shackman and Fox, 2016; Walker et al., 2009, 2003, Walker and Davis, 2008, 2002; Waraczynski, 2016). A dominant view has been that the BNST mediates the sustained (anxiety-like) fear responses to long-duration threats, including contexts (Hammack et al., 2015; Lee and Davis, 1997; Waddell et al., 2006; Walker and Davis, 1997). However, more recent data suggest that the BNST mediates conditioned fear to threat CSs (whether short or long in duration) that are poor predictors of when aversive outcomes occur (Daldrup et al., 2016; Goode et al., 2019; Goode and Maren, 2017; Lange et al., 2017). Consistent with this, we have recently reported that pharmacological inactivation of the BNST attenuates fear elicited by discrete auditory CSs that poorly predicted shock onset (Goode et al., 2019).

If temporal predictability, rather than stimulus modality or duration, is the critical factor determining BNST involvement in conditioned fear, then there should be factors in which contextual fear conditioning is independent of the BNST. Indeed, it has recently been reported that BNST lesions do not affect context fear conditioning when the footshock US occurs relatively soon (1 min) after an animal is placed in the conditioning context relative to those shocked after a long delay (10 min) (Hammack et al., 2015). However, in this study total exposure to the conditioning context was not equated, which therefore confounded the timing of shock onset (imminent or remote) with the duration of the context CS (short or long).

Here we sought to disentangle these factors by using a fear conditioning procedure that equated total context and shock exposure, while varying the placement-to-shock interval (1 min or 9 min). Animals experienced four 10-min contextual conditioning sessions in which they received a single shock per session that either occurred 1 min after the animal was placed in the conditioning context (“IMMINENT”) or 9 min after placement in the context (“REMOTE”). We hypothesized that the reversible inactivation of the BNST would impair the expression of contextual freezing in animals conditioned with temporally remote (9-min placement-to-shock), but not imminent (1-min placement to shock), USs. We also examined the context-dependence of conditioning by assessing the degree of context discrimination supported by the two conditioning procedures. Furthermore, we characterized behavioral features of each procedure, including shock-induced activity during conditioning, and the freezing latencies and bout durations of the rats during fear retrieval. Overall, we found that pharmacological disruption of the BNST was most effective in disrupting remote but not imminent shock.

## RESULTS

### Conditioning using imminent or remote shock is context-dependent

As a first step, we began by characterizing the topography and specificity of conditioned freezing produced by imminent or remote shock. Rats were placed in the conditioning context and received a footshock either 1 or 9 min after placement in the chamber (“IMMINENT” and “REMOTE” footshock, respectively); this procedure was conducted across four 10-min session. One day after the final conditioning session, animals underwent a counterbalanced test for context discrimination. A schematic of the behavioral design is shown in Figure 1. As shown in Figure 2, animals freezing behavior increased across each conditioning session for each conditioning procedure. ANOVAs of freezing during conditioning sessions I-IV revealed a main effect of time for each session (session I, repeated measures: *F*_9,252_ = 10.05, *p* < 0.0001; session II, repeated measures: *F*_9,252_ = 6.80, *p* < 0.0001; session III, repeated measures: *F*_9,252_ = 14.90, *p* < 0.0001; session IV, repeated measures: *F*_9,252_ = 20.88, *p* < 0.0001). Additionally, a main effect of conditioning procedure was detected for each session (session I: *F*_1,28_ = 32.02, *p* < 0.0001; session II: *F*_1,28_ = 79.42, *p* < 0.0001; session III: *F*_1,28_ = 25.41, *p* < 0.0001; session IV: *F*_1,28_ = 18.41, *p* < 0.0005). Significant time × conditioning procedure interactions were detected across conditioning (session I, repeated measures: *F*_9,252_ = 2.67, *p* < 0.01; session II, repeated measures: *F*_9,252_ = 8.21, *p* < 0.0001; session III, repeated measures: *F*_9,252_ = 12.09, *p* < 0.0001; session IV, repeated measures: *F*_9,252_ = 16.94, *p* < 0.0001). No main effects of test order or any other interactions were detected for any of the conditioning sessions (*F*’s < 2.70, *p*’s > 0.10). Nonetheless, it is worth noting that post-shock freezing greatly differed between the two procedures by the end of training. Specifically, a factorial ANOVA comparing freezing behavior of IMMINENT rats during minute 2 (session IV) vs. freezing of REMOTE rats during minute 10 (session IV) revealed a main effect of conditioning procedure (main effect of conditioning procedure: *F*_1,28_ = 86.33, *p* < 0.0001; no other main effects or interactions, *F*’s < 0.75, *p*’s > 0.40). Reduced post-shock freezing in REMOTE animals may be due to the short interval between shock and removal from the context (in this case, shock may signal removal from the chamber).

**Figure 1.**
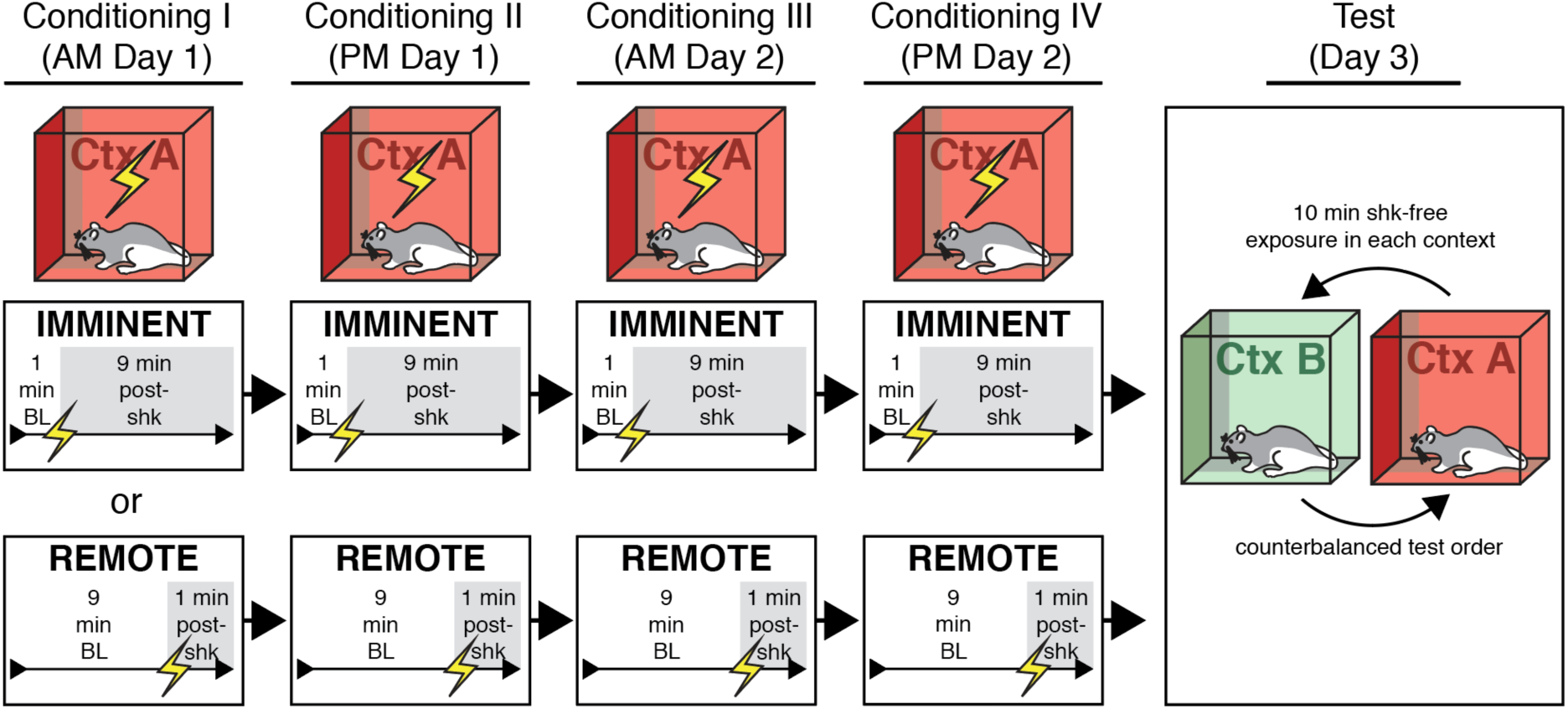
Behavioral design for testing context discrimination following conditioning using imminent and remote shock onset. Rats were randomly assigned to undergo four separate sessions of contextual fear conditioning using IMMINENT (1-min pre-shock baseline) or REMOTE (9-min pre-shock baseline) unsignaled shock. Each conditioning session was 10 min each. After training, and in a counterbalanced manner, IMMINENT and REMOTE rats were placed in a novel context (Context B) or the conditioning context (Context A) for 10 min in the absence of shock before being switched to the other context for an additional shock-free 10 min session.

**Figure 2.**
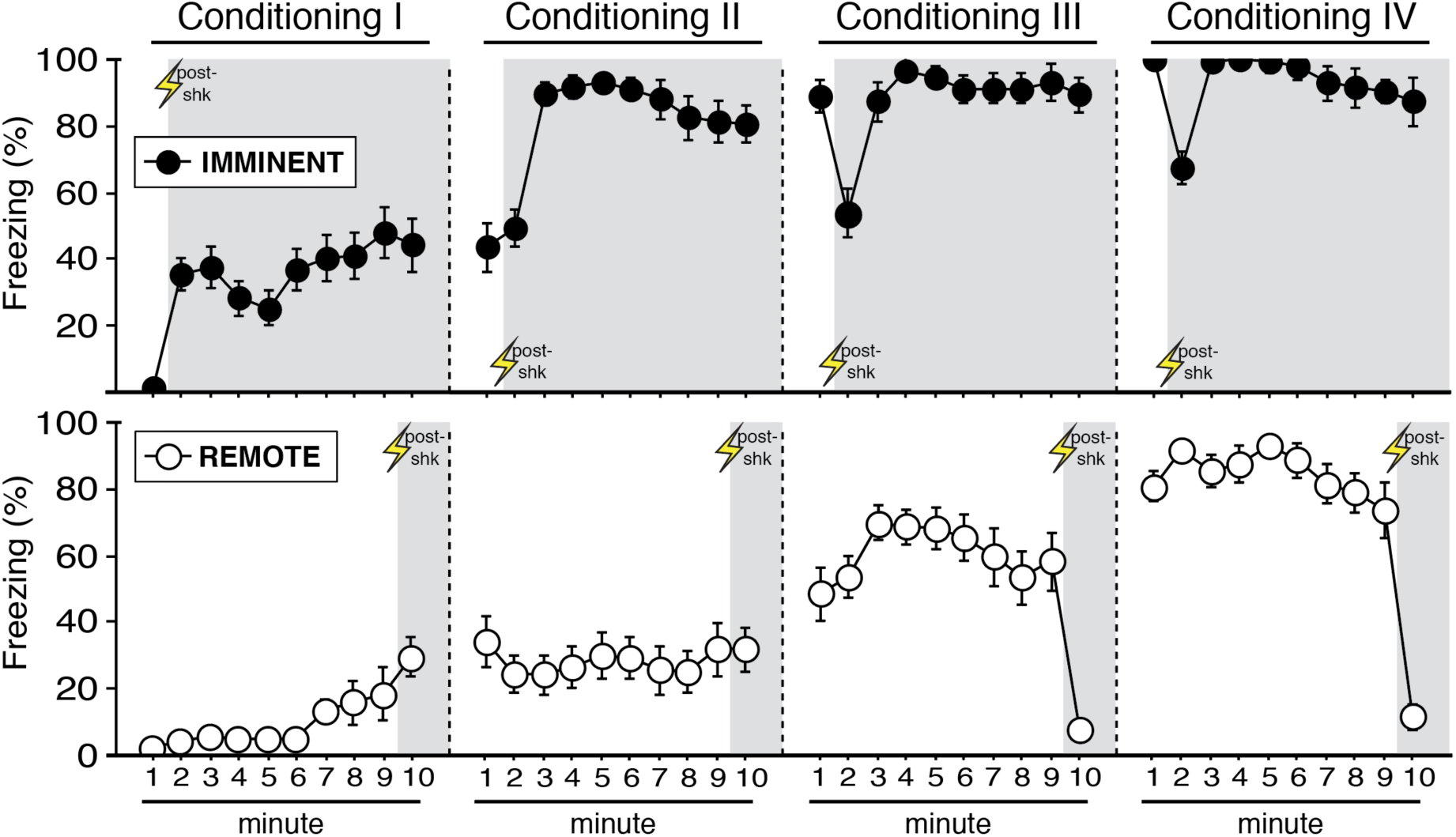
Context conditioning using imminent or remote shock onset. Mean percentage freezing (± s.e.m.) of IMMINENT (top) and REMOTE (bottom) rats at each minute of each conditioning session. Shaded areas indicate minutes post-shock for both procedures at each session. IMMINENT (*n* = 16); REMOTE (*n* = 16).

To test for retrieval and for the extent of context generalization, animals were either placed in the conditioning context or a novel context (10 min/session) followed by exposure to the alternate context ∼4 hrs later (test order was counterbalanced). As shown in Figure 3A and 3B, animals exhibited robust freezing in the conditioned context and lower fear in the novel context. Specifically, three-way ANOVA of freezing (with test context as a factor) revealed a main effect of context (*F*_1,56_ = 142.43, *p* < 0.0001), indicating robust context discrimination. Additionally, we observed a main effect of conditioning procedure (*F*_1,56_ = 7.38, *p* < 0.01), a main effect of time (repeated measures: *F*_9,504_ = 5.32, *p* < 0.0001), a time × context interaction (repeated measures: *F*_9,504_ = 3.30, *p* < 0.001), a time × conditioning procedure interaction (repeated measures: *F*_9,504_ = 17.85, *p* < 0.0001), a time × context × conditioning procedure interaction (repeated measures: *F*_9,504_ = 3.12, *p* < 0.005), and a time × context × test order interaction (repeated measures: *F*_9,504_ = 4.03, *p* < 0.0001). However, there was no overall main effect of test order (F < 0.09, p > 0.75). No other main effects or interactions were detected (*F*’s < 1.80, *p*’s > 0.06). A discrimination index (i.e., mean freezing of animals in the conditioning context subtracted from freezing percentages in the novel context) was calculated (Figure 3C), which revealed no significant difference in the extent of this discrimination between the two conditioning groups (*t* < 1.5, *p* > 0.15). In total, these data indicate that although both IMMINENT and REMOTE rats exhibited generalized freezing in the novel context, this generalized freezing was significantly less than freezing in the conditioned context. Additionally, the extent of context discrimination was similar between the training procedures, suggesting that both training procedures produce similar levels of context-dependent fear conditioning.

**Figure 3.**
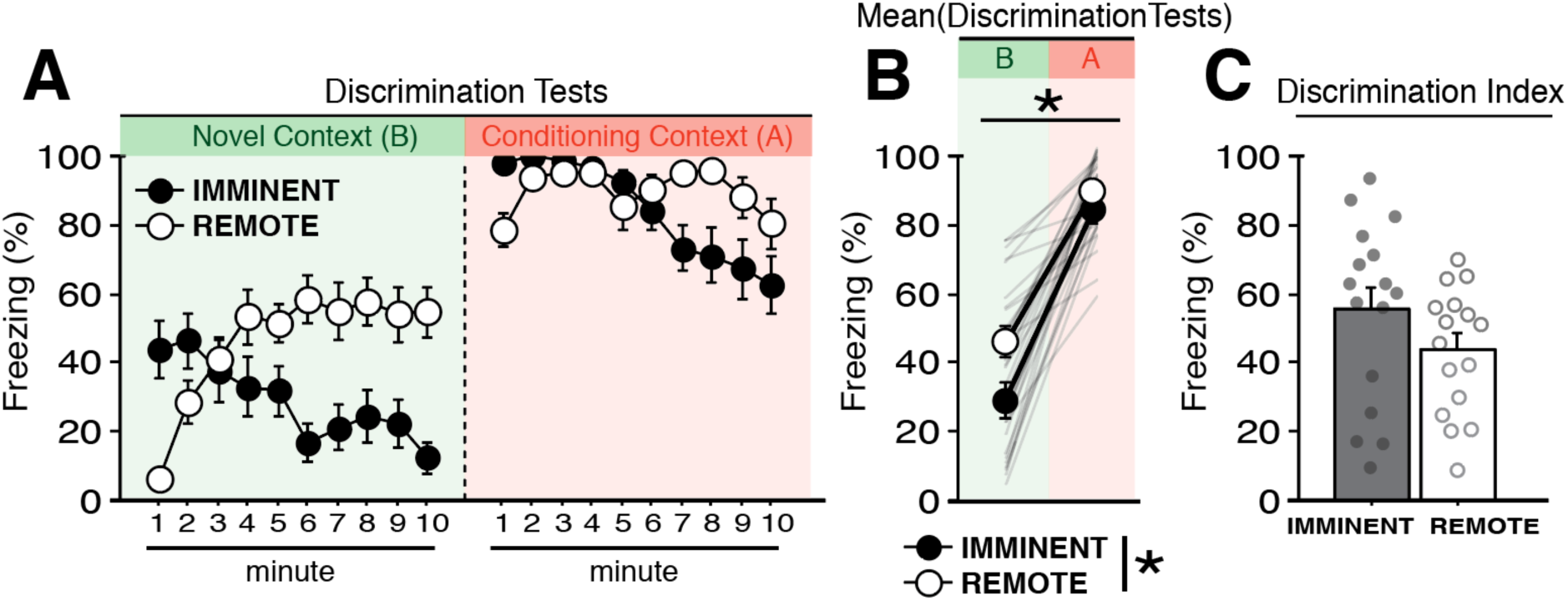
Context discrimination following imminent and remote shock training. **(A)** Mean percentage freezing (± s.e.m.) of IMMINENT and REMOTE rats at each minute of shock-free exposure to a novel context (Context B) or the conditioning context (Context A). **(B)** Mean percentage freezing (± s.e.m.) of IMMINENT and REMOTE rats across the entire test in each context. Shaded lines denote individual performance of each animal. **(C)** Mean freezing percentages (± s.e.m.) in Context B were subtracted from mean responding in Context A to generate a discrimination index for IMMINENT and REMOTE rats. IMMINENT (*n* = 16); REMOTE (*n* = 16). \**p* < 0.05.

### Conditioning procedure influences the topography of both activity-burst URs and freezing CRs

To determine whether the placement-to-shock interval influences the unconditioned response to footshock, we examined the magnitude of shock-evoked activity bursts (as assessed by cage displacement) during each 2-sec shock. Levels of shock-induced activity are shown in Figure 4A (left). Repeated measures ANOVA revealed a significant main effect of shock number (*F*_3,90_ = 7.25, *p* < 0.0005), indicating that shock-induced activity changed across the course of the conditioning sessions. A shock number × conditioning procedure interaction was detected in the ANOVA (*F*_3,90_ = 5.38, *p* < 0.005), indicating differences in the shock-induced activity between the two conditioning procedures across the conditioning sessions (no other main effects were detected: *F* < 0.5, *p* > 0.5). These differences are further apparent when comparing the percent change in shock-induced activity from the first to the final shock (Figure 4A; right). An unpaired *t*-test revealed that IMMINENT rats exhibited a significant reduction in the magnitude of the shock-induced activity across the trials as compared to REMOTE rats (*t*_30_ = −3.48, *p* < 0.005). These findings parallel our recent work (Goode et al., 2019), which found that conditioning-related decreases in shock-induced activity were lower in backward(unpredictable)-conditioned compared to forward (predictable)-conditioned animals. These outcomes may reflect a greater regulation of US responding in procedures in which animals can predict the onset of footshock.

**Figure 4.**
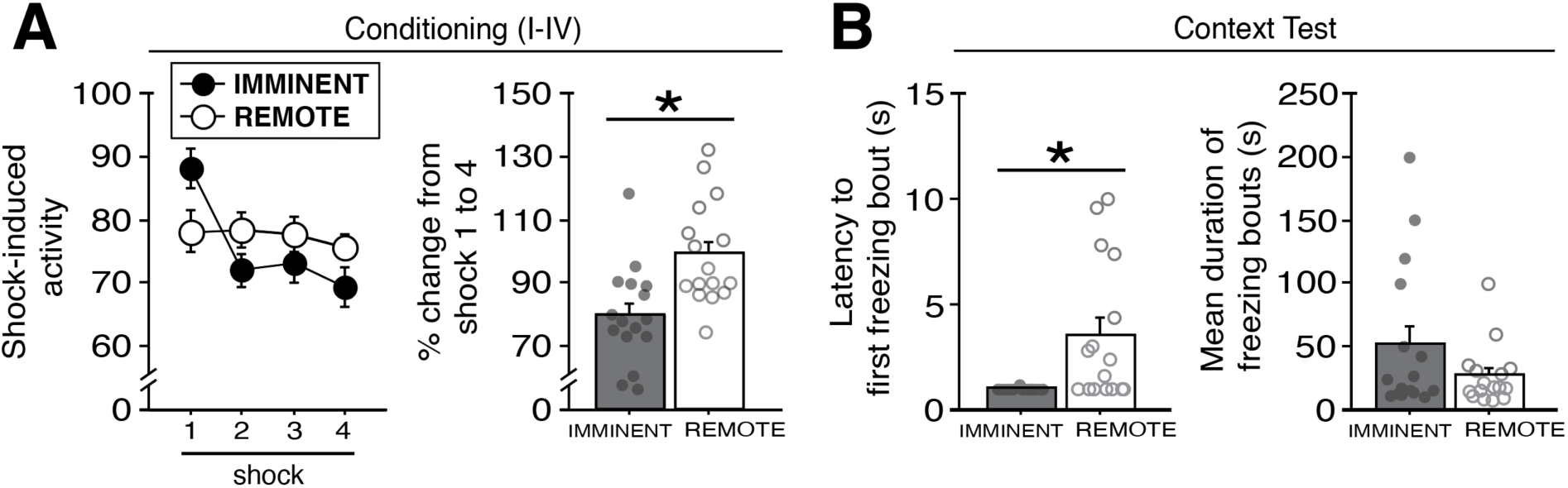
Shock-induced activity, latencies and durations of conditioned freezing bouts in animals trained with imminent or remote shock. **(A)** Left panel shows mean values of shock-induced activity (± s.e.m.) of IMMINENT and REMOTE rats at each shock (data correspond to the conditioning of rats shown in Figure 2). Right panel shows the percentage change in magnitude of shock-induced activity (± s.e.m.) from shock 1 to shock 4. **(B)** Left panel depicts latency (in seconds; ± s.e.m.) of IMMINENT and REMOTE rats to exhibit their first freezing bout in the conditioning context during testing (corresponding to Figure 3). Right panel shows mean duration of each bout (± s.e.m.) for the entire test. IMMINENT (*n* = 16); REMOTE (*n* = 16). \**p* < 0.05.

In addition to affecting the expression of shock-induced URs, the conditioning procedures produced differences in the nature of conditioned freezing in the conditioning context. To elucidate these differences, we examined the latency to the first freezing bout, as well as the average length of the freezing bouts in the conditioning context at test (Maren, 2001b). Although percentages of freezing are commonly reported as an index of learning, how animals reach certain magnitudes of freezing can differ. For example, an animal could express 50% freezing across a 10-min session by freezing for a sustained 300-sec bout, or by engaging in ten separate 30-sec bouts across the session. Thus, by examining latencies and durations of bouts over time, we may reveal important differences in BNST-dependent or -independent defensive strategies.

Latencies and bout durations of freezing of IMMINENT and REMOTE rats in the conditioned context are shown in Figure 4B (left). We opted to perform these analyses on the retrieval responses of IMMINENT and REMOTE rats from Figure 3 because these two groups exhibited similar overall magnitudes of freezing percentages across the 10-min session. Although both groups exhibited relatively short latencies for initiating freezing (all rats exhibited freezing within the first 15 sec of the exposure), IMMINENT rats were freezing almost immediately upon entering the context. Indeed, an unpaired *t*-test revealed that IMMINENT rats exhibited significantly shorter latencies to their first bout of freezing as compared to REMOTE rats (*t*_30_ = - 3.02, *p* < 0.01). Concerning the duration of the conditioned responses, we observed no significant difference between IMMINENT and REMOTE rats in the average length of freezing bouts (*t* < 1.5, *p* > 0.15). Thus, these data identified distinct (as well as overlapping) features of URs and CRs in REMOTE and IMMINENT rats, which may be factors the contribution of the BNST to the expression of contextual fear.

### Reversible inactivation of the BNST disrupts fear to contexts conditioned with remote, but not imminent, shock onset

In this experiment we tested whether placement-to-shock interval influences the role of the BNST in the expression of contextual freezing. Rats with cannula targeting the BNST were placed in the conditioning context and received a footshock either 1 or 9 min after placement in the chamber (“IMMINENT” and “REMOTE” footshock, respectively); a schematic of the behavioral design is shown in Figure 5. Cannula placements for all rats included in the analyses as well as a representative image of cannula tracts in tissue containing the BNST are displayed in Figure 5–Figure Supplement 1.

**Figure 5.**
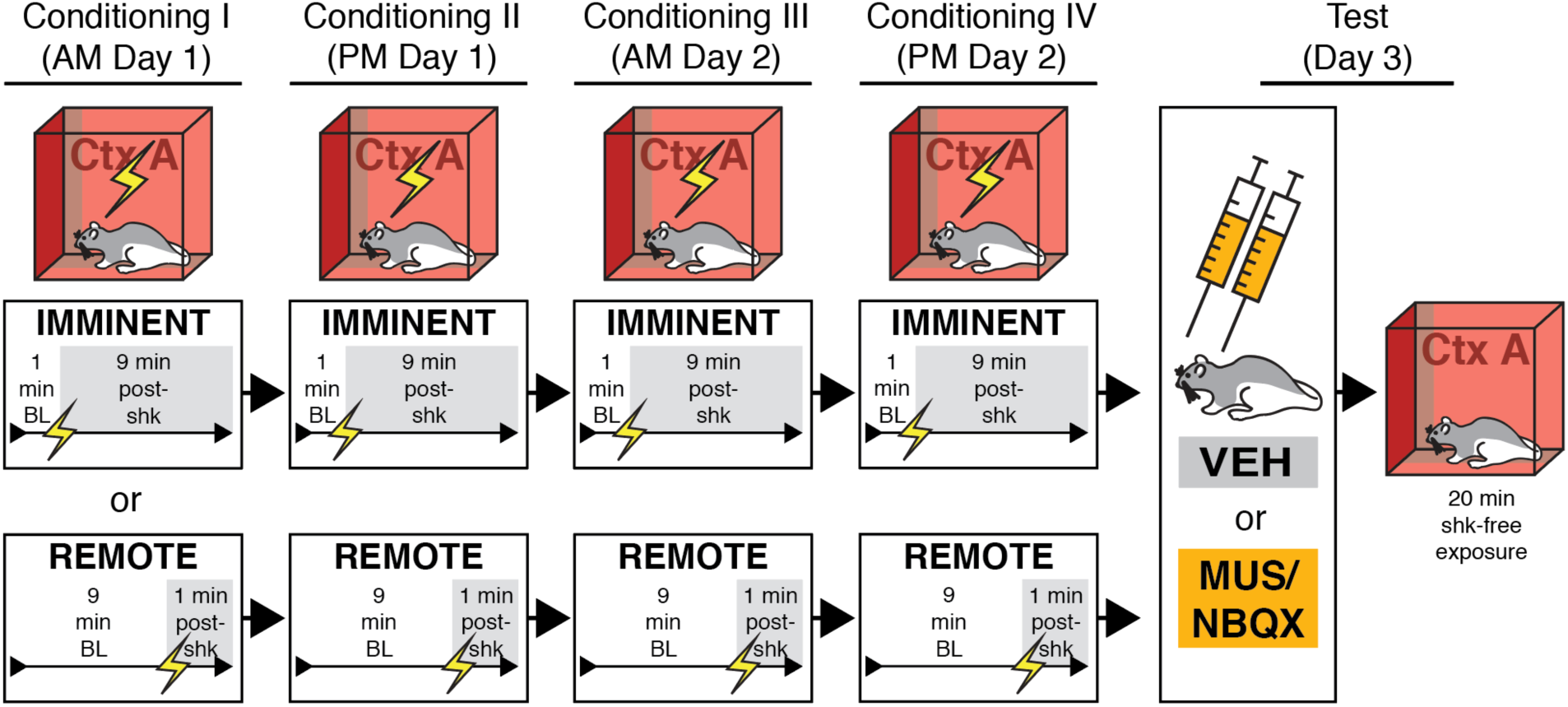
Behavioral design for contextual fear conditioning with imminent or remote footshock. BNST-cannulated animals were randomly assigned to undergo four separate sessions of contextual fear conditioning using IMMINENT (1-min pre-shock baseline) or REMOTE (9-min pre-shock baseline) unsignaled shock. Each conditioning session was 10 min. After conditioning, IMMINENT and REMOTE rats were infused with MUS/NBQX (to reversible inactivate) or vehicle (VEH) into the BNST prior to a 20 min shock-free retrieval session in the conditioning context.

Freezing during each minute of each of the four conditioning sessions is shown in Figure 6. Freezing behavior reliably increased across the conditioning sessions. Separate ANOVAs of freezing during each conditioning session (I-IV) revealed a main effect of time for each session (session I, repeated measures: *F*_9,396_ = 17.91, *p* < 0.0001; session II, repeated measures: *F*_9,396_ = 22.58, *p* < 0.0001; session III, repeated measures: *F*_9,396_ = 24.39, *p* < 0.0001; session IV, repeated measures: *F*_9,396_ = 17.17, *p* < 0.0001). Additionally, a main effect of conditioning procedure was detected for the first three sessions (session I: *F*_1,44_ = 14.62, *p* < 0.0005; session II: *F*_1,44_ = 50.11, *p* < 0.0001; session III: *F*_1,44_ = 27.85, *p* < 0.0001). Moreover, a time × conditioning procedure interaction was detected for each conditioning session (session I, repeated measures: *F*_9,396_ = 3.16, *p* < 0.005; session II, repeated measures: *F*_9,396_ = 11.17, *p* < 0.0001; session III, repeated measures: *F*_9,396_ = 6.46, *p* < 0.0001; session IV, repeated measures: *F*_9,396_ = 13.272, *p* < 0.0001). These data indicate that rats in the IMMINENT shock group generally increased their freezing more rapidly across each session.

**Figure 6.**
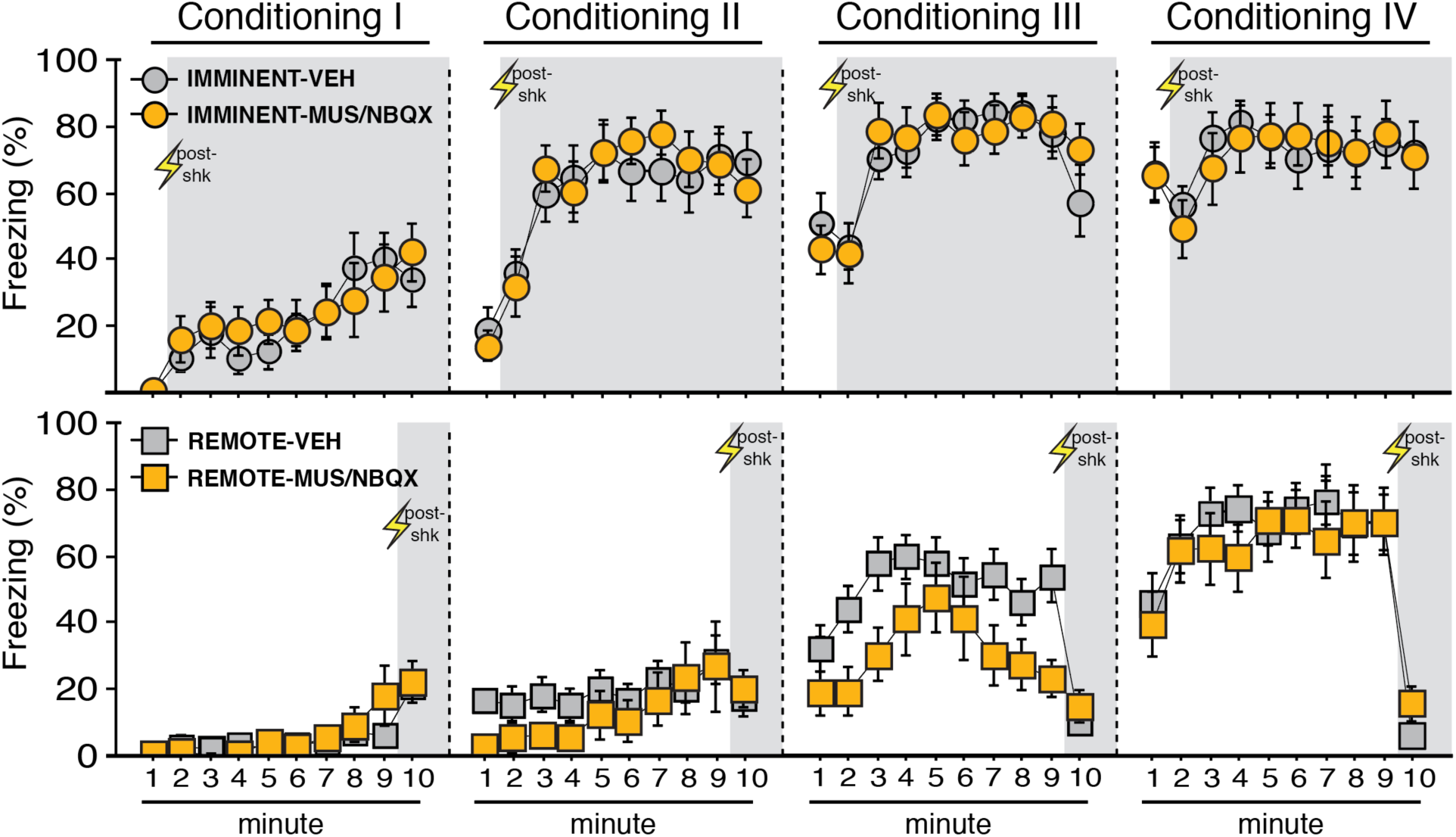
Context conditioning using IMMINENT or REMOTE shock onset. Mean percentage freezing (± s.e.m.) of IMMINENT (top) and REMOTE (bottom) rats at each minute of each conditioning session (I-IV). Shaded areas indicate minutes post-shock for both procedures at each session. Drug and vehicle assignments are shown for comparison but no BNST infusions occurred during this pre-test phase. IMMINENT-VEH (*n* = 13); IMMINENT-MUS/NBQX (*n* = 11); REMOTE-VEH (*n* = 15); REMOTE-MUS/NBQX (*n* = 9). \**p* < 0.05.

Rats that were slated to receive SAL or MUS during the test were similar to one another during conditioning (there were no drug infusions during conditioning). A time × drug assignment interaction was found for session 3 (repeated measures: *F*_9,396_ = 2.20, *p* < 0.05), but this difference was not apparent by session 4; no other main effects or interactions of drug assignment were found (*F*’s < 2.15, *p*’s > 0.15). Post-shock freezing in the final session appeared to mirror patterns seen in the prior experiment. Specifically, REMOTE-shocked animals exhibited significantly less freezing during minute 10 of session IV vs. minute 2 of session IV of IMMINENT animals (factorial ANOVA, main effect of conditioning procedure: *F*_1,44_ = 59.387, *p* < 0.0001; no other main effects or interactions, *F*’s < 2.25, *p*’s > 0.14). Collectively, these data reveal reliable increases in freezing during conditioning; results that are similar to the prior experiment.

After conditioning, the GABA_A_ receptor agonist, MUS, or the AMPA receptor antagonist, NBQX (collapsed into a single group, “MUS/NBQX”; see methods), was used to inactivate the BNST prior to a shock-free retrieval test in the conditioning context (Figure 7). A significant main effect of time (repeated measures: *F*_19,836_ = 7.02, *p* < 0.0001) and a significant time × conditioning procedure interaction (repeated measures: *F*_19,836_ = 1.90, *p* < 0.05) were found in the ANOVA, indicating that IMMINENT and REMOTE rats broadly changed in freezing across the course of the test. A main effect of drug was also apparent (*F*_1,44_ = 5.215, *p* < 0.05). A significant time × conditioning procedure × drug assignment interaction was also found in the ANOVA (repeated measures: *F*_19,836_ = 1.81, *p* < 0.05). Fisher’s PLSD indicated REMOTE-MUS/NBQX exhibited significantly less freezing as compared to REMOTE-VEH (*p* < 0.01). No other main effects or interactions were detected in the analyses (*F*’s < 3.60, *p*’s > 0.05). Neither drug on its own was effective in attenuating freezing in IMMINENT animals (NBQX vs. VEH: *F*’s < 0.45, *p*’s > 0.90; MUS vs. VEH: *F*’s < 0.70, *p*’s > 0.85]. In total, these data indicate that contextual fear expression after conditioning with imminent shock is insensitive to inactivation of the BNST.

**Figure 7.**
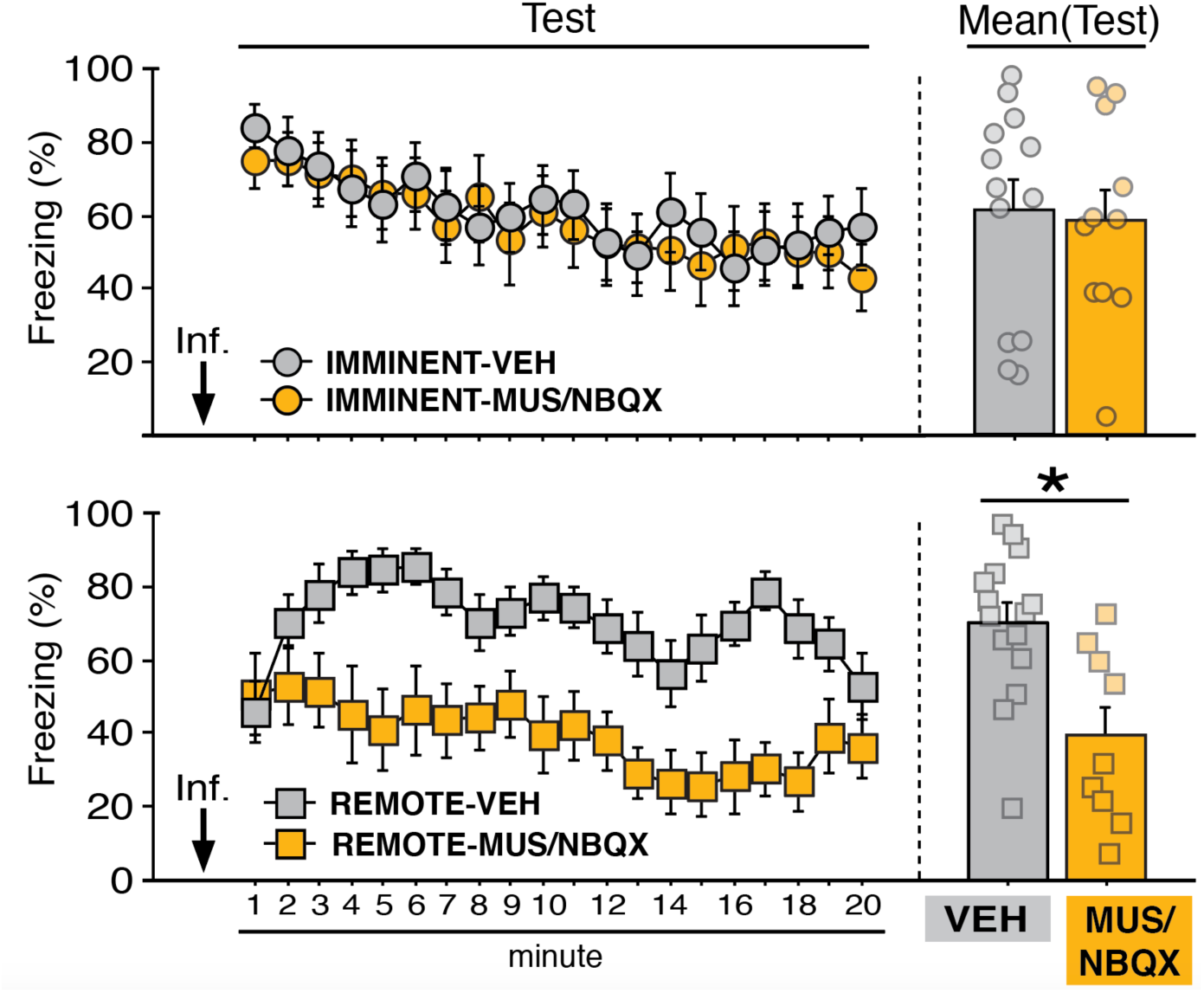
BNST inactivation attenuates freezing in a context conditioned with remote but not imminent shock onset. Mean percentage freezing (± s.e.m.) of IMMINENT (top) and REMOTE (bottom) rats in the conditioning context following intra-BNST microinfusions of VEH or MUS/NBQX. IMMINENT-VEH (*n* = 13); IMMINENT-MUS/NBQX (*n* = 11); REMOTE-VEH (*n* = 15); REMOTE-MUS/NBQX (*n* = 9). \**p* < 0.05.

## DISCUSSION

We demonstrate a dissociable role for the BNST in the expression of fear to contexts that signal imminent vs. remote shock onset. Inactivation of the BNST impaired conditioned freezing in a context associated with remote footshock, but had no effect on freezing in a context associated with imminent footshock. Importantly, the conditioning procedures equated total context and shock exposure and produced similar degrees of context-dependent freezing, revealing that it is the timing of shock with respect to placement in the conditioning context that determines involvement of the BNST in contextual freezing. Interestingly, despite differences in latency to freeze, the topography of freezing behavior was similar in animals conditioned with imminent and remote footshocks. This reveals that although freezing was similarly sustained across the context retrieval test, BNST inactivation only reduced freezing in animals conditioned with remote footshock. This suggests that it is not the duration of the CR that determines BNST involvement in conditioned freezing, but rather the degree to which the context or CS signals when footshock will occur.

The current results build on a prior study from our lab (Goode et al., 2019), in which we found that CSs that signaled imminent shock (e.g., forward-trained CSs) were insensitive to BNST inactivation, whereas CSs that were poor predictors of shock onset (e.g., backward or randomized CSs) were sensitive to the manipulation. Importantly, the current work replicates the findings of a significant prior study that showed that context fear was insensitive to lesions of the BNST when trained with early shock onset (Hammack et al., 2015). The current study builds on and expands on these findings in several critical ways. First, the study by Hammack and colleagues (2015) utilized permanent excitotoxic lesions that persisted throughout conditioning and retrieval, making it difficult to isolate whether the role of the BNST in context fear was specific to processes of conditioning, consolidation, or retrieval. Although the BNST may have roles during these other stages, our current data suggest that the BNST is necessary at retrieval for proper recall following remote- but not imminent-shock training. Additionally, Hammack and colleagues (2015) compared context fear expression in two groups of animals that not only differed in the timing of shock, but also in total context exposure. In the current study, all animals had equal exposure to the context, indicating that the effects on retrieval are precisely because of shock timing, rather than the duration of the context exposure *per se*.

Prior work has suggested that the BNST mediates distinct forms of fear expression, particularly sustained responses, that are dissociable from phasic fear responses, which may be governed by other structures such as the central amygdala [e.g., (Sullivan et al., 2004; Walker and Davis, 1997)]. In contrast, we observed robust freezing in both IMMINENT and REMOTE animals, including similar overall durations of freezing bouts, but only REMOTE training was sensitive to BNST inactivation. Thus, we believe a more accurate depiction of BNST function is that it mediates responses to remote and unpredictable threats, and that these responses may in some cases be sustained (perhaps as the risk of threat persists), but that response duration is not always predictive of whether BNST is involved.

We also observed differences in the magnitude of shock-evoked URs of IMMINENT vs. REMOTE animals across the conditioning sessions. Specifically, imminent-shock onset appeared to coincide with reductions in shock-induced activity across the sessions, whereas remote-shocked animals largely expressed consistent levels of activity-burst URs across the sessions. We recently reported a similar outcome in the US-induced activity of animals subjected to auditory forward vs. backward fear conditioning (Goode et al., 2019), insofar as forward-conditioned animals more rapidly exhibited reductions in their activity across conditioning. These outcomes may reflect a greater regulation of US responding in signaled or imminent threat paradigms, which may be a factor in BNST recruitment to fear.

Overall, our findings concord with recent human imaging studies examining unpredictability [e.g., (Clauss et al., 2019; Figel et al., 2019; Naaz et al., 2019)], which may support a role for the BNST in anticipatory responses to temporally unpredictable threatening stimuli. Unpredictability of the onset of aversive events may serve as a common thread for the BNST’s broad contributions to anxiety, as well as for fear and drug relapse the aftermath of unpredictable stressors (Goode et al., 2018; Goode and Maren, 2019; Harris and Winder, 2018; Mantsch et al., 2016; Miles et al., 2018; Stamatakis et al., 2014; Vranjkovic et al., 2017). Nonetheless, the BNST is an intricate, sexually dimorphic and heterogeneous structure, with diverse functions (Avery et al., 2014; Crestani et al., 2013; Daniel and Rainnie, 2016; Flavin and Winder, 2013; Hammack et al., 2012, 2010, 2009; Kash, 2012; Kash et al., 2015; Radley and Johnson, 2018; Waraczynski, 2016)—more work is needed to fully characterize its complex neural responses and contributions to aversive stimuli (Acca et al., 2017; Duvarci et al., 2009; Haufler et al., 2013; Luyck et al., 2018, 2017; Martinon et al., 2019; Moaddab and Dabrowska, 2017). To conclude, we build on our prior study (Goode et al., 2019) by demonstrating a dissociable role of the BNST in the behavioral expression of contextual fear, an effect that depends on the timing of shock onset.

## MATERIALS AND METHODS

### Subjects

All subjects were adult (200-240 g) male Long-Evans (Blue Spruce) rats (*n* = 96, before exclusions) obtained from Envigo (Indianapolis, IN). Rats were individually housed in a climate-controlled vivarium and kept on a fixed light/dark cycle (lights on from 7:00 AM to 9:00 PM). Home cages consisted of a clear plastic cage layered with sawdust bedding (changed weekly), with access for the animals to standard rodent chow and water *ad libitum*. Home cages were housed on a rotating cage rack. Group assignments for all behavioral testing was randomized for cage position on the racks. Prior to the start of any surgical or behavioral procedures, experimenters handled the rats (∼30 sec/day) for five consecutive days. All procedures were performed in accordance with the US National Institutes of Health (NIH) Guide for the Care and Use of Laboratory Animals and were approved by the Texas A&M University Institutional Animal Care and Use Committee.

### Apparatuses

Conditioning/testing chambers (30 cm × 24 cm × 21 cm; MED Associates, Inc.) were housed in two distinct rooms of the laboratory (eight chambers per room). Each chamber rested in larger external sound-attenuating cabinets. The chambers were comprised of aluminum (sidewalls) and Plexiglas (rear walls, ceilings, and front doors). The floors of the chambers consisted of a grid of nineteen stainless steel bars (4 mm in diameter), spaced 1.5 cm apart (center to center). For delivery of footshock, the grid floors were connected to an electric shock source and a solid-state grid scrambler (MED Associates, Inc.). The chambers were also equipped with 15 W house lights (as needed for distinct contexts). Fans in each cabinet provided background noise (∼70 dB) as needed. Aluminum pans collected animal waste below each chamber. Speakers were attached to the chambers (for delivery of auditory tones), but these were not used in the current study.

To measure freezing over time, our lab utilized an automated and unbiased scoring system. Specifically, each chamber rests on a load-cell platform that detects chamber displacement as the animal moves. Load-cell activity values (in a range of −10 to +10 V) were digitized at 5 Hz and recorded using Threshold Activity Software (MED Associates, Inc.). These measurements are converted offline to generate absolute values ranging from 0 to 100; low values correspond to minimal chamber displacement. Thus, freezing bouts were set as absolute values of ≤10 for 1 s or more. Percentages of freezing can then be calculated for periods of time as defined in each experiment. For shock-induced activity measurements, we reported the absolute values generated by the Threshold Activity Software, such that larger values indicate more displacement of the chamber as a result of the animal’s activity (Goode et al., 2019; Maren, 1998).

Experiments utilized distinct contextual features to generate two different contexts. Each context was assigned to a separate behavioral room in the laboratory. Chambers were cleaned with the context’s respective odor before and after each squad of rats. For Context A, the following procedures and features were used: Testing chambers were wiped down with 3% acetic acid odor, and a small amount of the solution was poured into the pans beneath the chambers. Chamber lights were turned on, while the cabinet fans remained off. The cupboard doors of the cabinets were closed. A dim red light was used for illuminating the room. Rats were transported to and from the context using small white plastic boxes. For Context B, the following procedures were used: 1% ammonium hydroxide odor was used to scent the chambers. Chamber lights were turned off, while the cabinet fans remained on. Thin black plastic sheets were set on top of the grid floors. The cupboard doors of the cabinet were left open. White lights were used to illuminate the behavioral room. Rats were transported to and from Context B using sawdust-containing black plastic transport boxes.

### Surgeries

For the data corresponding to Figs. 1-3, animals were first implanted with bilateral guide aimed at the BNST [similar to prior reports: (Acca et al., 2017; Goode et al., 2019, 2015; Nagaya et al., 2015; Zimmerman and Maren, 2011)]. On the day of surgery, animals were individually transported from the vivarium to a surgical suite and prepped for surgery. Animals were deeply anesthetized using gaseous isoflurane (5% for induction; maintained during surgery at 1-2%). Once deeply anesthetized, animals were secured in a stereotactic frame (Kopf Instruments, Inc.), the hair on the top of the head was clipped, artificial tear ointment was applied, and the skin at the site of the incision was treated with povidone-iodine. A small incision was made in the skin and the skull exposed. Small holes were drilled into the skull to attach jeweler’s screws. Bregma and lambda of the skull were aligned on an even plane and small holes were drilled in the skull to allow for insertion of bilateral stainless-steel guide cannulas (26-gauge, 8 mm from the bottom of their plastic pedestals; Small Parts). The guide cannulas were lowered into the brain at the following coordinates: −0.15 mm posterior to bregma, ±2.65 mm lateral to the midline, and −5.85 mm dorsal to dura (guide cannulas were angled at 10**°** with their needles directed at the midline). Dental cement was applied to cover the skull and to secure the guide cannulas to the screws. Stainless steel obturators (33-gauge, 9 mm; Small Parts) were inserted into the guide cannulas. Subsequently, rats were removed from the stereotaxic frame, topical antibiotic ointment was applied to the head, and the rats were monitored for recovery. Rats were provided rimadyl-containing bacon-flavored tablets for post-operative pain management. Animals recovered for one week in their homecages prior to the onset of behavioral training.

### Intracranial infusions

In the week of recovery following surgery, animals were acclimated to the process of intracranial microinfusions. This involved transporting the animals (in sawdust-containing five-gallon buckets) from the vivarium (in squads of eight) to the separate room used for the infusions in the laboratory. The stainless-steel obturators were gently removed from the guide cannulas and replaced with clean ones. Animals were then returned to their homecages. This process was repeated twice on separate days.

For the data shown in Figures 5-7, the γ-aminobutyric acid (GABA)_A_ receptor agonist, muscimol (MUS), or the α-amino-3-hydroxy-5-methyl-4-isoxazolepropionic acid (AMPA) receptor antagonist 2,3-dihydroxy-6-nitro-7-sulfamoyl-benzo[f]quinoxaline-2,3-dione, NBQX, was used to temporarily inactivate the BNST. Both MUS (Bangasser et al., 2005; Breitfeld et al., 2015; Buffalari and See, 2011; Fendt et al., 2003; Goode et al., 2019, 2015; Markham et al., 2009; Pina et al., 2015; Sajdyk et al., 2008; Xu et al., 2012) and NBQX (Adami et al., 2017; Davis and Walker, 2014; Goode et al., 2019, 2015; Zimmerman and Maren, 2011) have been used to reversibly inactivate the BNST. MUS (Sigma-Aldrich) was dissolved in physiological saline to a concentration of 0.1 μg/μl. NBQX (Sigma Life Sciences) was dissolved in physiological saline to a concentration of 10.0 μg/μl. Physiological saline served as vehicle (“VEH” for all cases). An equal number of animals were assigned to received muscimol (“MUS”; *n* = 16 prior to exclusions) or NBQX (*n* = 16 prior to exclusions). Ultimately, MUS- (*n* = 8, after exclusions; included in figures) and NBQX-treated (*n* = 12, after exclusions; included in figures) animals were collapsed into a single group (“MUS/NBQX”) as neither manipulation alone altered responding to the early-shock context (see Results for additional details).

For the purpose of microinjections, 9 mm stainless steel injectors (33 gauge, Small Parts) were connected to water-filled polyethylene tubing (PE-20; Braintree Scientific), with the tubing’s other end connected to gastight 10 μl syringes (Hamilton, Co.). Syringes were secured to an infusion pump (KD Scientific, Inc.). The infusion pump was set to deliver a total volume of 0.275 ul (per injector) of MUS, NBQX, or VEH; infused at a rate of 0.275 μl/min. On the day of infusions, MUS, NBQX, or VEH was drawn up into the injectors and tubing, with a small air bubble separating the drugs or vehicle from the water in the tubing. Rats (in squads of eight) were brought to the infusion room and the obturators were removed. The drug- and vehicle-filled injectors were inserted into the guide cannulas; microinjections occurred simultaneously for the entire squad of rats (injectors were left in the cannulas for 1 min following the infusions to allow for diffusion). Once the infusions were completed, the injectors were removed and new obturators were inserted. Animals were then immediately transported to the behavioral chambers as necessary for testing.

### Behavioral procedures and exclusions

Summaries of the behavioral procedures can be found in Figs. 1 and 4. For all experiments, footshock (2 sec, 1.0 mA) served as the unconditioned stimulus (US).

*IMMINENT/REMOTE w/ contextual discrimination.* In a 2 × 2 design, rats (*n* = 32) were randomly selected to experience context conditioning using early (“IMMINENT”) or late (“REMOTE”) onset of shock, and to undergo a counterbalanced test order (Context A then B, or Context B then A) for the context discrimination test. No rats were excluded from these analyses, resulting in the following final group sizes (as shown in the figures): IMMINENT (*n* = 16); REMOTE (*n* =16). If collapsed to account for test order (i.e., IMMINENT w/ test order one, IMMINENT w/ test order two, REMOTE w/ test order one, and REMOTE w/ test order two), then each group is comprised of eight rats. On the morning of the first day of behavioral training, animals (in squads of eight rats) were transported from the vivarium to Context A. We alternated squads undergoing IMMINENT or REMOTE training. Drug and vehicle assignments were counterbalanced across each squad and for chamber position during training and testing. For rats undergoing IMMINENT training, animals were placed in the chamber and allowed to acclimate to the context for 1 min before the onset of the US; animals remained in the chamber for 9 min after shock onset before being removed and returned to their homecages (session I). Later that afternoon, this process was repeated (session II). The following day, this process was repeated for a morning session (III) and an afternoon session (IV). For REMOTE rats, the animals were placed in Context A and allowed to acclimate for 9 min before the onset of the US; rats remained in the chamber for 1 min after shock onset before being returned to their homecages (session I). This process was repeated for an afternoon session (II), and two more session, III and IV, on the following morning and afternoon, respectively. Thus, both IMMINENT and REMOTE rats experienced four conditioning sessions in total (two per day; each conditioning session was 10 min). On the day after the final conditioning session, rats (in squads comprised of equal numbers of IMMINENT and REMOTE animals) were placed in either the novel (Context B) or conditioning (Context A) context for 10 min in the absence of shock. We alternated squads for novel or conditioning context exposure until each rat had experienced both the novel and conditioning context for 10 min each. Both of these tests occurred on the same day (day three of behavioral procedures). After each test session, animals were returned to their homecages until sacrificed at a later date.

*IMMINENT/REMOTE w/ BNST inactivation.* In a 2 × 2 design, rats (*n* = 64, prior to exclusions) were randomly selected to experience contextual conditioning using early (“IMMINENT”) or late (“REMOTE”) onset of shock, and to undergo drug (“MUS/NBQX”) or vehicle (“VEH”) infusions into the BNST prior to testing. Of the original sixty-four rats, fifteen were found to have off-target cannulas and were excluded from the final analyses. An additional rat was euthanized to due illness and was excluded from the study. This resulted in the final group sizes for the final analyses (as shown in the figures): IMMINENT-VEH (*n* = 13); IMMINENT-MUS/NBQX (*n* = 11; comprised of five MUS-treated and six NBQX-treated animals); REMOTE-VEH (*n* = 15); REMOTE-MUS/NBQX (*n* = 9; comprised of seven MUS-treated and two NBQX-treated animals). For contextual fear conditioning, IMMINENT and REMOTE rats experienced identical conditioning procedures as discussed above, with drug assignments being counterbalanced for each squad as possible for training. After completing the final conditioning session, and on the following day, animals were infused with MUS/NBQX or VEH immediately before a 20-min shock-free retrieval test in the conditioning context. In this case, test squads included equal numbers of IMMINENT and REMOTE rats (counterbalanced for drug assignments). After the test session, animals were returned to their homecages.

### Histological procedures and image analyses

For animals implanted with cannulas in the BNST, and at the conclusion of behavioral procedures, rats were overdosed on sodium pentobarbital (Fatal Plus; 100 mg/ml, 0.5 ml, i.p.) and transcardially perfused using chilled physiological saline followed by 10% formalin. Brains were extracted and placed in 10% formalin for 24 hrs at 4° C. Brains were then switched to a 30% sucrose-formalin solution until sectioning occurred (stored at 4° C). For sectioning, brains were flash frozen using crushed dry ice and coronal sections (40 μm) containing the BNST were collected using a cryostat set to −20° C (Leica Microsystems). Sections were wet-mounted onto gelatin-subbed glass microscope slides. Subsequently, the tissue was stained with 0.25% thionin using a standard staining procedure. Glass coverslips were glued (Permount, Sigma) to the microscope slides, and the slides were allowed to dry before imaging. Photomicrographs of the thionin-stained tissue were generated at 10× using a Leica Microscope (MZFLIII) and Leica Firecam software. Data shown in Figs. 5-7 include only those animals with injector tips localized to within the borders of the BNST.

### Statistics

All data were submitted to repeated or factorial analysis of variance (ANOVA) or two tailed *t*-tests as described for each experiment. Only after a significant omnibus *F* ratio in the ANOVA (α was set at 0.05) were data submitted to post-hoc comparisons in the form of Fisher’s protected least significant difference (PLSD). All data are shown as means (± s.e.m). No statistical methods were used to predetermine group sizes (group sizes were selected as based on prior work). Data distributions were assumed to be normal, although these were not formally tested.

## ACKNOWLEDGEMENTS

The authors thank Carolyn Evemy and Sohmee Kim for technical assistance. Supported by grants from the National Institutes of Health (R01MH065961 and R01MH117852 to S.M. and F31MH107113 to T.D.G.), as well as a McKnight Foundation Memory and Cognitive Disorders Award and a Brain & Behavior Research Foundation NARSAD Distinguished Investigator Grant to S.M.

---

**Figure 5-Figure Supplement 1.**
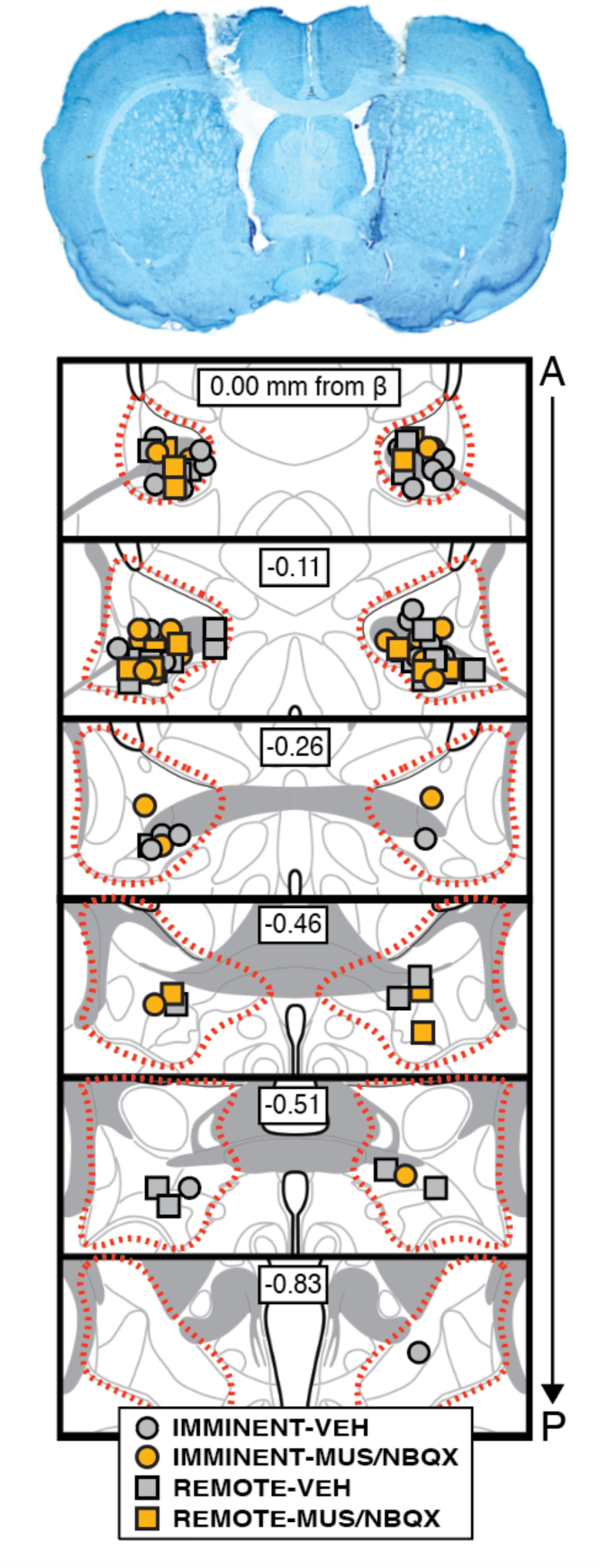
Bilateral cannula placements in the BNST. Representative photomicrograph of a thionin-stained coronal section (40 μm) depicting bilateral cannula tracts and injector tips in the BNST (top image). Bottom image shows the positions of injector tips for each animal for each group included in the final analyses of the experiment (approximate borders of the BNST are outline by the dotted red line). IMMINENT-VEH (*n* = 13); IMMINENT-MUS/NBQX (*n* = 11); REMOTE-VEH (*n* = 15); REMOTE-MUS/NBQX (*n* = 9).

